# Neural Representational Geometry of Feature Binding Operations

**DOI:** 10.64898/2026.02.18.706604

**Authors:** Laura Sainz Villalba, P. Michael Furlong, Madeleine Bartlett, Nicole Sandra-Yaffa Dumont

**Affiliations:** Institute of Neuroinformatics, ETH Zurich and University of Zurich; University of Waterloo Collaboration Centre, National Research Council of Canada

**Keywords:** neural representation geometry, feature binding, vector symbolic algebra, neurosymbolic programming

## Abstract

The brain faces the feature binding problem: how are multiple stimulus features and variables combined into coherent representations that support flexible behavior? A key finding from neuroscience is that some brain regions employ factorized representations, where distinct features are encoded in neural state space in such a way that enables independent readout and robust generalization. Various algebraic operations have been proposed to model multi-variable representations, but despite extensive study of their theoretical properties (e.g., capacity, noise robustness), it remains unclear which operations produce the representational geometries observed in neural recordings. We systematically evaluate six binding operations implemented in recurrent spiking neural networks performing a working memory task. We find that only superposition and binding with slot-filler structure produce factorized geometry with favorable scaling, while the alternatives do not. These results provide a taxonomy linking algebraic binding operations to neural representational signatures, offering guidance for both computational modelers and experimentalists.

## Introduction

Representations of features and variables from distinct brain circuits must combine to form coherent wholes on which we learn, act, and remember. This challenge is known as the feature binding problem. Understanding how the brain solves this problem requires examining how multiple features are jointly represented in neural populations. Recent work on the representational geometry of such activity has revealed that the geometry of co-encoded representations varies across brain regions and task contexts (Bernardi et al., 2020; Boyle et al., 2024; Courellis et al., 2024; Ebitz & Hayden, 2021; Gao et al., 2017; Low et al., 2018).

Of particular interest are *factorized* representations, in which different variables are encoded in an (abstract) way that supports generalization across conditions (Bernardi et al., 2020). Such representations have been observed in hippocampus, dorsolateral prefrontal cortex, and anterior cingulate cortex (Bernardi et al., 2020; Mishchanchuk et al., 2024). Factorized geometry develops with learning, appears in subjects performing inferential reasoning, and correlates with task performance and structure complexity (Courellis et al., 2024; Sainz Villalba et al., 2025).

These analyses have revealed much, but they alone do not explain the computational mechanisms underlying binding.

Several proposals model conjoined representations via algebraic combinations of high-dimensional vectors (including Tensor Product Representations (Smolensky, 1990) and Vector Symbolic Algebras (Kleyko et al., 2022; Schlegel et al., 2022)) which can be represented via neural activity with population codes (Thomas et al., 2021). While these operations have been evaluated for capacity and noise robustness (Carzaniga et al., 2025; Clarkson et al., 2023; Kleyko et al., 2022), they have not been subjected to a neuroscience-inspired analysis of their representational geometry.

We address this gap by systematically comparing six binding operations (including concatenation, tensor product, summation, and convolution; see Fig. 1) implemented in recurrent spiking neural networks. Our analysis reveals which mechanisms achieve factorized geometry, characterizes trade-offs between factorization and discrimination capacity, and provides principled guidance for modeling binding in specific brain regions or cognitive functions.

**Figure 1.**
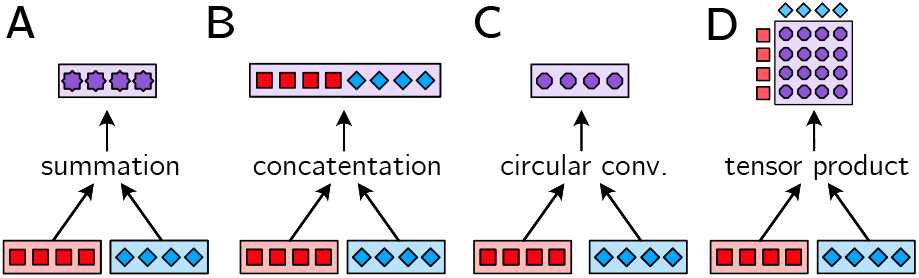
Illustration of binding operations.

**Figure 2.**
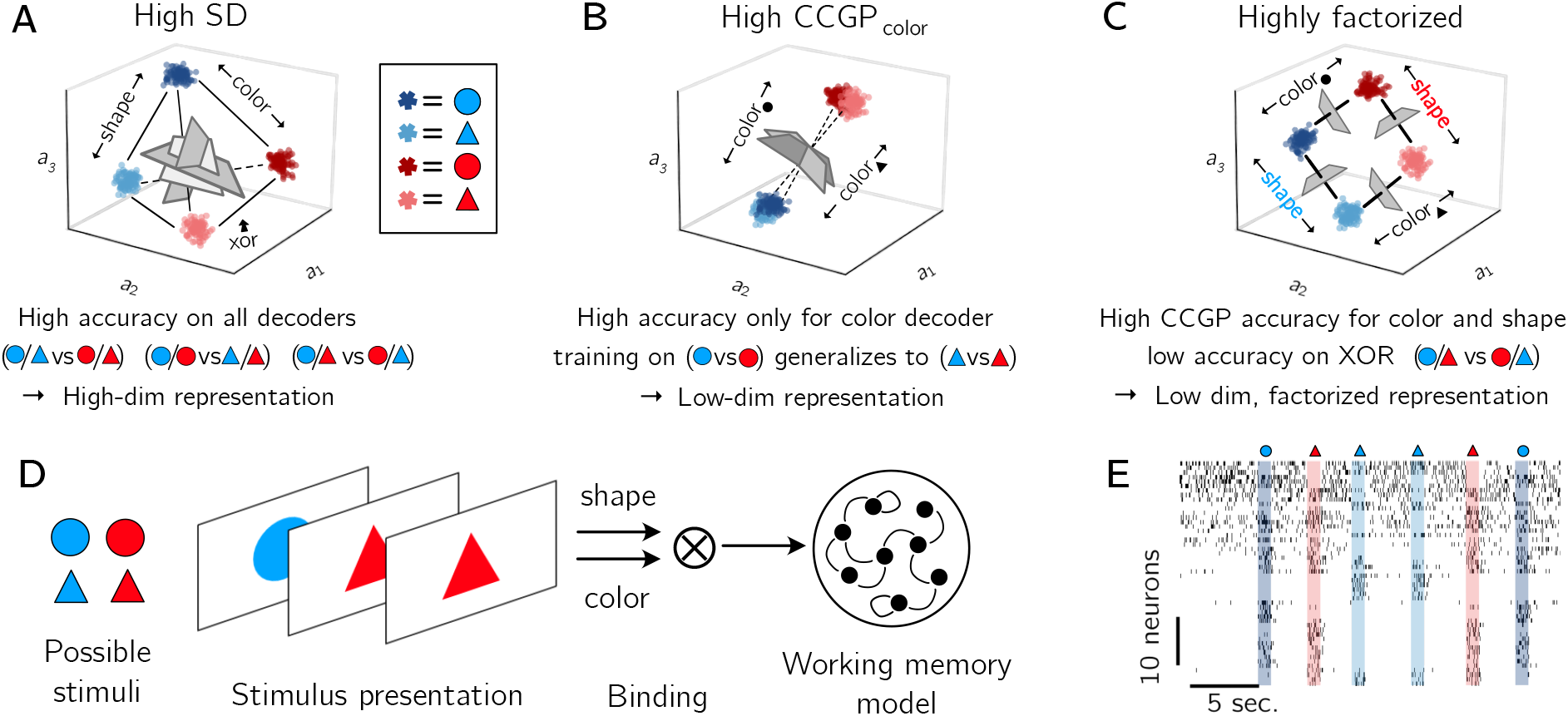
Overall of different neural representation geometries and our experimental setup. Here we are considering a neural population representing two variables simultaneously (shape and color), each of which can take on one of two values (circle/triangle and blue/red). **(A)** A high shattering dimensionality (SD) means that one can linearly decode many dichotomies (binary parsings on conditions) from the activity. Even a random encoding in a high enough dimensionality space will have a high SD. **(B)** A high CCGP means that a linear decoder trained in one condition (the value of a different variable) generalizes to other conditions. High CCGP for one variable does not imply high CCGP in another. **(C)** A factorized representation has high CCGP for all variables of interest. However, if such a representation is low dimensional then the XOR dichotomy will not be decodable. In the extreme case, dimensionality for a factorized representation would correspond with the number of variables. **(D)** In this paper, we simulate a recurrent neural population. In variable-length trial blocks, different combinations of shape-color variables are presented for 1s. Vector representations of these variables are bound with a mathematical operation and injected into the recurrent population during the presentation period. The recurrent weights are set so that the population is a short term working memory. **(E)** A spike raster plot of the recorded activity over the experiment with presentation periods highlighted and labeled. We use activity in different conditions to compute linear decoders.

### Related Concepts

The concept of factorized representations connects to several ideas across fields, though terminology varies. A variable is encoded in *abstract* format (Bernardi et al., 2020) if it can be decoded robustly regardless of other variables. *Cross-condition generalization performance* (CCGP) quantifies this by measuring how well a linear decoder generalizes to held-out conditions. A conjoined representation is *factorized* when its constituent variables each exhibit high CCGP against one another—that is, the decoding axis for each variable remains approximately parallel across values of other variables, enabling generalization (Bernardi et al., 2020), (see Fig.2A-C). A related concept in cognitive science is *compositional* representations, which refers to complex mental structures built by combining simpler constituents, e.g., ‘‘red square’’ combines a color and shape. Factorized neural representations support compositionality: if color and shape are encoded such that they can be independently decoded and recombined, systematic generalization becomes possible.

In neuroscience, neural population activity can be viewed as a point in a high-dimensional state space, where each neuron contributes one dimension. Population codes are described as *orthogonalized* when activity patterns for different experimental conditions lie along approximately orthogonal axes in this space. However, orthogonality alone is not sufficient for factorization as the feature dimensions themselves must be orthogonal and compositional, not just particular feature combinations. And vice-versa, orthogonality of decoding dimensions is not strictly required to ensure a factorized representation of them. In machine learning, *disentangled* representations are individual latent dimensions that correspond to independent factors of variation in the data (Burgess et al., 2018; Locatello et al., 2019), and are mathematically equivalent to factorized representations.

## Methods

We study how different binding operations shape neural representations. We consider joint encoding of two object features: color and shape. These features are represented as high-dimensional vectors, which are combined using binding operations from prior theoretical proposals. The resulting bound representations are maintained in a recurrent spiking neural network performing a working memory task, and analyzed as simulated neural recordings. By holding the task, feature statistics, memory dynamics, and analysis methods constant while varying only the binding operation, we isolate how different binding schemes give rise to distinct representational geometries and decoding properties at the population level.

### Binding Operations

The binding problem can be formalized as follows: given two feature vectors, **f**_1_, **f**_2_ ∈ℝ^*d*^, representing, for example, COLOR=RED and SHAPE=CIRCLE, we can produce a bound representation **b** ∈ℝ^*d*^′that (1) preserves information about both features, (2) can be inverted to recover the constituent features, and (3) supports compositional operations. Different binding mechanisms make different tradeoffs in terms of *d*′ (the dimensionality of the bound representation), the fidelity of feature recovery, and the resulting representational geometry.

The simplest binding operation is *concatenation*: **b** = [**f**_1_; **f**_2_] ∈ℝ^2*d*^ (*e*.*g*. Soll et al., 2019). This approach trivially achieves perfect factorization, as features occupy non-overlapping subspaces, but dimensionality grows linearly with the number of bound features. It is computationally expensive and biologically implausible for binding many features. Smolensky proposed using the binding of a ‘‘slot’’ of a feature vector with the value ‘‘filling’’ that slot, feature binding can then be represented by summing these slot-filler pairs (Smolensky, 1990) . This method exploits the pseudo-orthogonality of random high-dimensional vectors to let multiple variables exist in one vector representation. More recently, direct tensor products between different features, 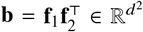, have been used to represent multi-modal data in neural networks (Zadeh et al., 2017; Zhao et al., 2024). While tensor products are an expressive way to represent bound variables, the resulting representation has dimensionality *d*^2^, scaling more poorly than concatenation.

Vector symbolic algebras (VSAs) provide dimensionality-preserving alternatives to tensor binding (Kleyko et al., 2022; Levy & Gayler, 2008; Plate, 1995). In VSAs, feature vectors can be combined by *superposition* (typically, vector addition), which produces new vectors that retain similarity with its constituent elements, and *binding*, which produces new vectors that are dissimilar to their constituent elements. The binding operator is integral to defining a VSA, and many different operators have been proposed (reviewed in Schlegel et al., 2022), including:

- **Circular convolution**: element-wise multiplication in the Fourier domain, as used in Holographic Reduced Representations (Plate, 1995)
- **Vector-derived transformation binding (VTB)**: permutation-based binding (Gosmann & Eliasmith, 2019)
- Binding for binary or bipolar vectors: element-wise products, XOR

VSA operators can be implemented in spiking neural networks (Eliasmith, 2013; T. C.Stewart et al., 2011) and have been used successfully to model various cognitive tasks (Agerskov, 2016; Dumont et al., 2022; Furlong & Eliasmith, 2024; Renner et al., 2022; T. Stewart et al., 2012). Often, both superposition and these binding operators are used to create hierarchical, structured representations. For example, a slot-filler structure using circular convolution represents a red circle as **b** = **s**_color_ ⊗ **f**_red_ + **s**_shape_ ⊗ **f**_circle_ ∈ ℝ^*d*^ .

### Binding Simulation

#### Binding operations

We test the following operations to combine features/variables: vector sum, concatenation, tensor product, VTB, circular convolution, and a slot-filler structure using circular convolution. The bound representation produced by each operation is then provided as input to a recurrent working memory network. This network is a leaky integrator of the vector-valued network inputs that maintains activity (stimulus presentation lasts 1 s, see Fig.2).

#### Simulating neural recording

Neural activity is simulated using the Neural Engineering Framework (NEF) as implemented in the Nengo simulator (Eliasmith & Anderson, 2003). In this framework, a vector-valued signal **b** ∈ℝ^*d*^ is represented by the activity of a population of spiking neurons. Each neuron receives an input current given by

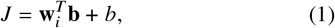

where **w**_*i*_ is the *i*^th^ neuron’s synaptic weights,corresponding to the neuron’s preferred direction in the represented space, and *b* is a bias current. The **w**_*i*_ are random vectors drawn from a uniform distribution over the unit hypersphere. This results in a distributed, heterogeneous population code for **b**.

Neurons are modeled as noisy leaky integrate-and-fire (LIF) units with standard membrane dynamics. Spiking activity is generated by thresholding the membrane potential, followed by a reset and refractory period. Population activity thus provides a noisy, time-varying representation of the underlying vector signal, analogous to extracellular recordings from biological neural populations.^1^

#### Population decoding analysis

For the analysis, we first binned neural activity into 200ms windows, with an overlap of 20 ms to smooth bin transitions, following similar procedures of neural data analysis applied to biological recordings (Boyle et al., 2024; Sainz Villalba et al., 2025). We then aligned neuron activities to the stimulus onset, and trained and tested individual decoders (linear support vector machines, Pedregosa et al., 2011) at each time step. For cross-validation, we randomly divided data into two non-overlapping groups of trials, with a train-test split of 80-20 % and averaged accuracy over 100 iterations. To avoid confounding factors during decoding of each variable, trials corresponding to each trial type combination (according to our 2×2 variable design) were split independently. Then, we randomly sampled population vectors from the corresponding trials in the training or testing set. Trials were resampled to ensure balanced conditions with a feature matrix of dimensions 2*n* trial samples by *n* features (number of neurons), with *n* being the number of sub-sampled neurons fixed to 80% unless otherwise specified.

#### Cross decoding and shattering dimensionality

We used CCGP, as described in Bernardi et al. (2020) and applied it to every time step. CCGP quantifies the performance (decoding accuracy) of a decoder trained to distinguish between values of a certain variable (main variable) and tested in a different set of trials that differ in another variable (condition). For example, to test generalized encoding of color, the decoder was trained to discriminate blue vs red stimuli in triangle-shaped trials and its performance was tested on decoding color in circle-shaped trials and vise versa. Then the CCGP for color is the average across all possible ways to split the train and test set (dichotomy) according to the rest of the conditions of interest, in this case according to shape. Factorized format for variables *i* and *j* corresponds to high CCGP decoding accuracy for both, invariant against each other. To compute Shattering Dimensionality (SD), we parsed our variable value combinations (red-triangles, blue-triangles, red-circles, blue-circles) in all possible (balanced) dichotomies (shape, color and XOR). The XOR dichotomy is (red-circle, blue-triangle) vs. (red-triangle, blue-circle). We then computed the average decoding accuracy across all, by training and testing (in disjoint trial sets), assuming the assigned binary parsing as the ground truth for labels. SD then relates to the capacity of encoding different condition combinations (parsings) as distinct, reflecting the intrinsic dimensionality of the neural representation (number of latent variables to explain population activity) (Fig.2). The embedding dimensionality (ED) (number of linearly independent variables to explain variability) was computed with principal component analysis (PCA) at each timestep across trials, subsampling 80% of neurons for 100 separate iterations. The minimal number of components for which the variance explained converged (< 1% marginal increase) was used as the ED result.

## Results

We computed linear decoding accuracy and CCGP at every timestep using 512 neurons; results for a single simulation of three operators are shown in Fig. 3. We found that 50 neurons were sufficient for high decoding accuracy, so we used this subset for subsequent analyses. Scores (CCGP for both shape and color, and XOR, the remaining dichotomy) were averaged over the stimulus presentation period and across 10 random seeds (Fig. 4).

**Figure 3.**
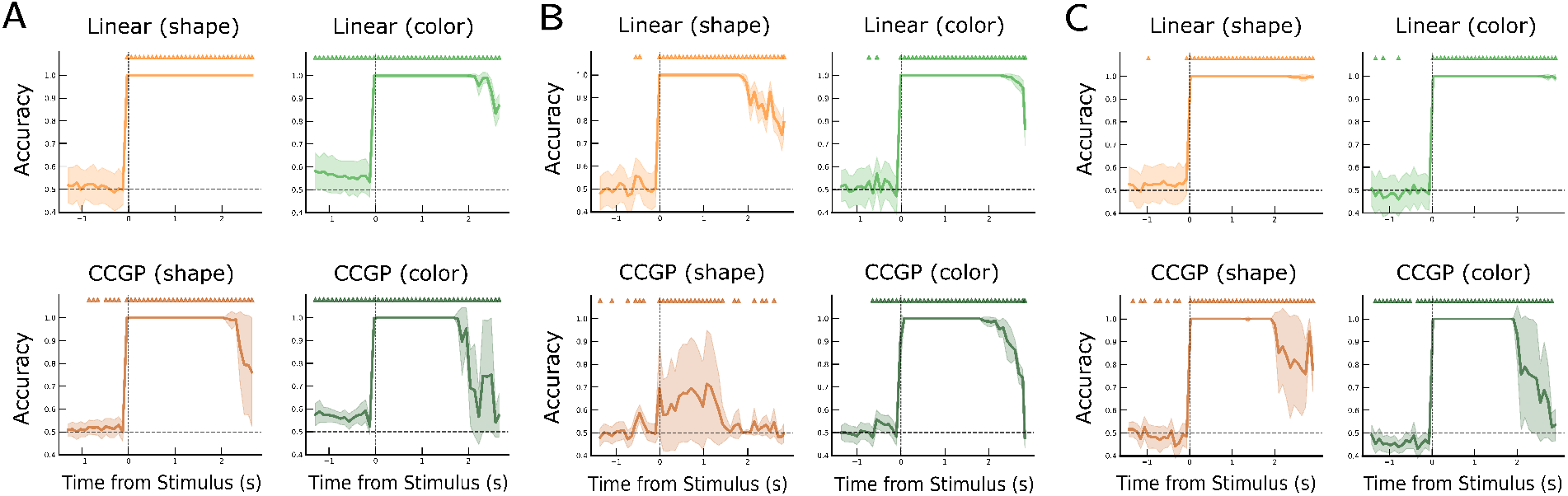
Timeseries for linear and cross-decoding of both shape and color across different binding operations: **(A)** Concatenation, **(B)** circular convolution, and **(C)** slot-filler, computed with an architecture of 512 neurons. Note that concatenation trivially expresses high CCGP values for both variables reflecting orthogonal subspaces. However, feature binding scales with *n d*. Circular convolution only expresses high CCGP for one variable, in this example for color, which may be the dominating decoding axis in the representation. Slot-filler is consistently able to encode both variables in a factorized format. Triangles denote significant timesteps (p-value < 0.05, t-test for independent samples with Bonferroni correction for multiple comparisons across timepoints).

**Figure 4.**
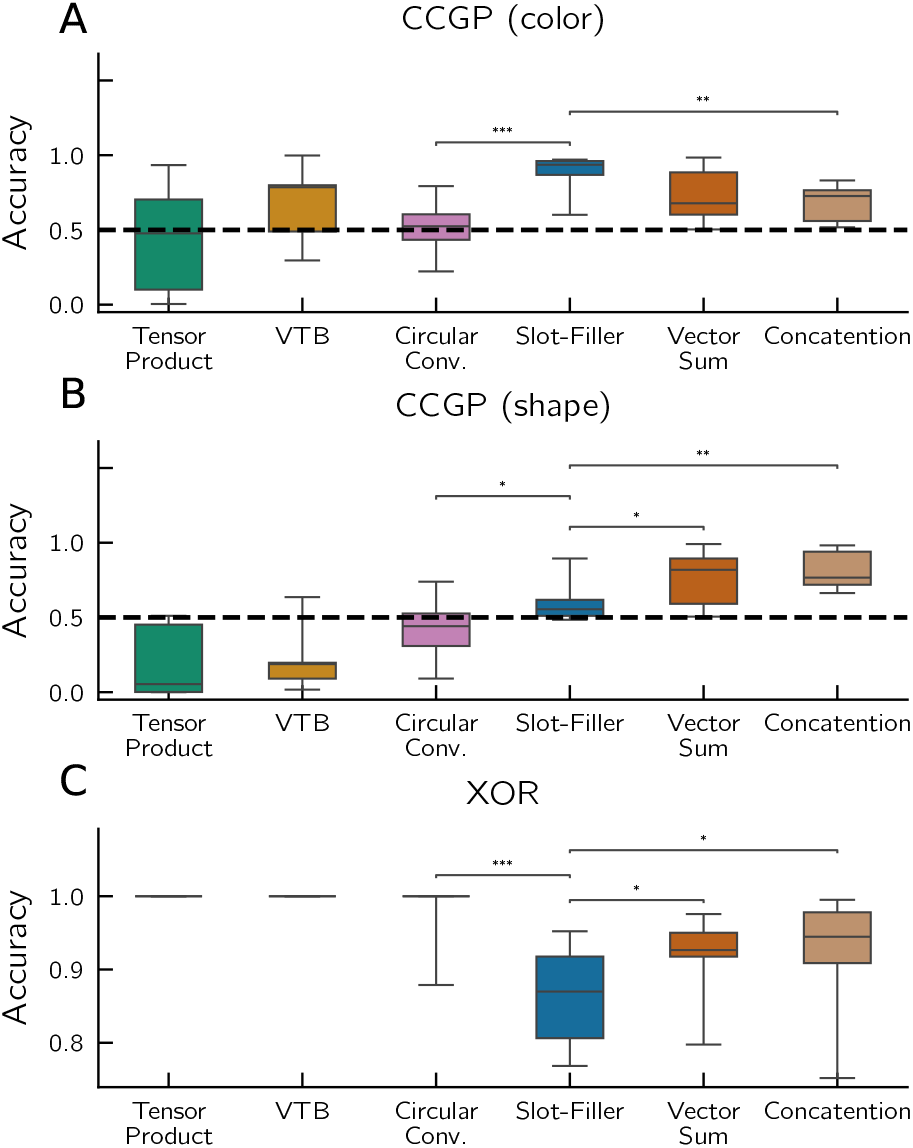
The CCGP and XOR performance for the different binding operations. We compute the average score over the full post-stimulus onset period. The boxes show the interquartile range of these scores over 10 seeds (a random initialization of variable vectors and network hyper-parameters). **(A-B)** The CCGP for (A) color and (B) shape. The dashed line at 0.5 marks chance performance. The binding operators are ordered from lowest to highest average CCGP. **(C)** The XOR-decoding performance. (Mann Whitney test, uncorrected. p-value: *: <0.05, **:0.01, ***:0.001)

All binding operations supported accurate retrieval of individual feature values (Fig.3, first row). However, only sum, concatenation, and slot-filler variants exhibited high CCGP for both color and shape, indicating factorized representations (Figs. 3, second row, and 4). Other operators failed to robustly encode both variables in abstract format but did have high XOR decoding accuracy. For example, circular convolution shows high CCGP for only one variable in 3B and CCGP near chance when averaged over many seeds (Fig.4).

The slot-filler operation, despite achieving reasonable mean XOR accuracy (>0.8), was significantly lower than sum or concatenation (Fig. 4C). This suggests slot-filler produces a more strongly factorized representation, approaching a lower-dimensional manifold at the cost of reduced capacity to enable XOR parsing (like the example in Fig.2C).

Supporting the results of high XOR accuracy, the SD was maximal (1, accuracy average across all parsing dichotomies) during stimulus presentation (Fig.5A). Similar SD timeseries were found for all binding operations. Importantly, the increase in SD did not correspond to an increase in ED but rather a drastic reduction from 30 during the pre-stimulus period to 3 during stimulus presentation. This reflects the structured activity of the population in response to the stimulus, arranging in a geometry that still maximally preserves the representation capacity with the minimal dimensionality.

**Figure 5.**
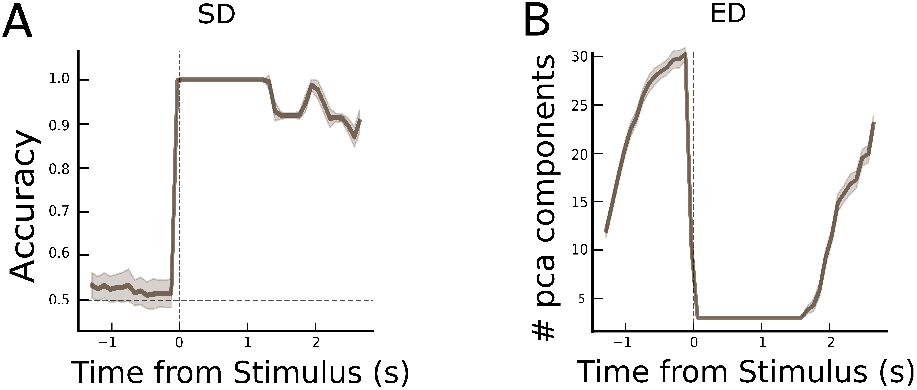
Shattering dimensionality (SD) and embedding dimensionality (ED) timeseries. The dynamics observed in this example (concatenation) reproduce across tested binding operations. After stimulus onset, population activity lies on a low-dimensional manifold explained with 3 components as compared to the pre-stimulus period where population activity is unstructured. This effect, however, does not prevent of being able to decode all condition dichotomies but rather expresses high SD reflecting structured collective activity

These dimensionality dynamics (SD and ED) do not ensure a factorized representation. This is evident when projecting neural activity during stimulus presentation into a 3D space via PCA and plotting decoding hyperplanes (Fig. 6). For slot-filler binding (Fig. 6A), the hyperplane for decoding shape trained on blue trials is approximately aligned with the hyperplane trained on red trials – in other words, shape is encoded abstractly. In contrast, with circular convolution (Fig.6B), these hyperplanes are approximately orthogonal. Despite both binding operations exhibiting similar dimensionality dynamics, only the former arranges trial conditions in a way that supports robust, disentangled readout of both variables (i.e., high CCGP).

**Figure 6.**
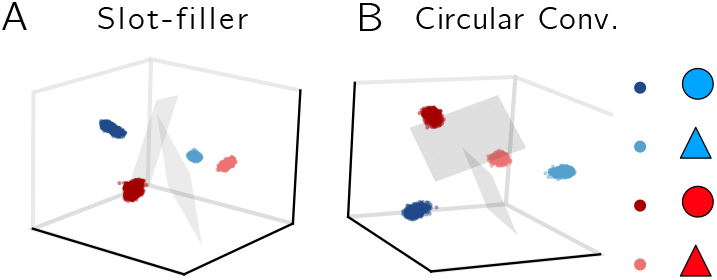
Neural activity during all stimulus presentations projected into a 3D space via PCA. Each point corresponds to activity at a given time, colored by input stimulus at that time. The gray planes are the shape separating hyperplanes for the two color contexts. The gray planes correspond to decoding hyperplanes for shape across the two possible different color contexts. For the slot-filler **(A)** we find that the hyperplanes are almost parallel approaching the collinear aligned hyperplanes found in perfectly factorized representations (see Fig.2C). In circular convolution **(B)**, by contrast, hyperplanes are orthogonal precluding generalization for shape across color.

## Discussion

### Factorized versus entangled

In this work, we analyzed the representational geometries of different algebraic binding operations. We subjected six binding operators to analysis techniques developed for neural data analysis. All operations, unsurprisingly, encoded both intended variables with high accuracy. However, only some (sum, concatenation, and slot-filler^2^) expressed conjoined representations in a factorized format as defined in the neuroscience literature. Of these, concatenation has limited biological plausibility for more than two features since it scales poorly. The remaining operations (sum and slot-filler) that do have more appropriate scalable properties(𝒪 (*d*))depart slightly in the trade-off between generalization capabilities (CCGP) and representation capacity (indirectly from XOR dichotomy) (see Fig.4).

These results can inform modeling choices by helping researchers select operators that match the factorization and dimensionality observed in a target brain region, ensuring that algebraic models of cognition better reflect biological computation.

Conversely, when neuroscientists observe particular representational geometries, these results suggest which neural architectures might produce them and which computations (e.g., generalization) would be enabled or precluded, offering support for hypotheses about underlying function. One key consideration in such modeling is the trade-off between generalization and discrimination, as noted in Boyle et al. (2024). A representation with high SD but chance CCGP encodes variables in an entangled manner. While this hinders compositional generalization, it may better support memory and learning in contexts where distinctiveness matters. Consider learning a function over bound representations: if representations are factorized, values learned in one context generalize to others, but this generalization can also cause interference and impede learning when contexts should be kept separate.

### Disambiguating variables in superposition

Both summation and slot-filler have good scaling and exhibited factorized geometry. However, slot-filler offers additional structure that avoids ambiguity. Consider representing multiple objects with their colors: a simple sum (**b** = **c**_1_+**c**_2_) conflates which color belongs to which object, while slot-filler (**b** = **s**_1_ ® **c**_1_ + **s**_2_ ® **c**_2_) preserves this correspondence. Furthermore, circular convolution is approximately invertible, enabling a network to compute 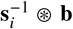to selectively retrieve a specific bound variables. Switching the slot effectively switches which variable is accessed – a mechanism that could interface naturally with attention.

### Limitations of linear decoding

The linear decoding techniques used to characterize factorization in neuroscience may be blind to more sophisticated representations. For example, it is sometimes useful to represent both individual features in superposition and their conjunction (**b**_2_ = **f**_1_ + **f**_2_+ **f**_1_ ⊗ **f**_2_). Through linear decoding, **b**_1_ and **b**_2_ may appear indistinguishable, yet they support different computations (both generalization and discrimination). This motivates building biologically plausible models that can be evaluated on both representational geometry and behavior, ensuring that matches to neural data are not merely superficial.

### Learning factorized representations

We have assumed a mature system with already-learned factorized representations. However, experimental work shows that factorized geometry exists contextually (Boyle et al., 2024) and can emerge through learning (Sainz Villalba et al., 2025). The mechanisms underlying this remain unclear: it could arise from constraints on standard learning rules, from interfacing circuits that shape learning, or top-down, as suggested by the induction of abstract representations via language instructions (Courellis et al., 2024).

This opens avenues for theoretical and experimental work on learning conjunctive representation. Notably, in the VSA literature, the problem of learning how to combine features (which should be summed, bound, nested, etc.) remains unsolved. Insights from neuroscience on how factorized representations emerge could inform this open problem.

### Scaling to more features and continuous variables

Our analysis considered two discrete features. Expanding to systems with more variables would test scalability and efficiency. VSAs also offer methods for representing continuous variables such as location or size, which have been applied in hippocampal models (Dumont et al., 2022). Simulations with algebraic binding operations in more complex tasks could generate predictions for experimental work studying multi-feature and continuous representations.

## Conclusion

We provide the first systematic comparison of algebraic binding operations evaluated through the lens of neural representational geometry. By applying analysis techniques from neuroscience (CCGP and SD) to operations proposed in cognitive science and VSA literature, we establish which mechanisms produce the factorized geometry observed in areas such as the hippocampus, dorsolateral prefrontal cortex, and anterior cingulate cortex.

Our results yield a clear taxonomy: sum and slot-filler binding produce factorized representations with favorable scaling, while circular convolution, VTB, and tensor product do not. These operations do not result in factorization because bound representations are dissimilar to their constituents, enabling discrimination but precluding generalization. Between the factorized operations, slot-filler offers additional structure that preserves variable-feature correspondence and supports selective retrieval.

This work bridges terminology across fields. Concepts such as ‘orthogonalized’, ‘disentangled’, ‘factorized’, and ‘compositional representations’ have developed somewhat independently in neuroscience, machine learning, and cognitive science. We clarify their relationships and dependencies, identifying which properties are strictly required for particular computational capabilities and which emerge as byproducts.

Our taxonomy offers practical guidance: modelers can select binding operations that match the geometry observed in target brain regions, while neuroscientists can use observed geometry to constrain hypotheses about underlying circuit mechanisms and the computations they enable. A key open question is whether the learning mechanisms that produce factorized representations in biological systems inherently favor certain binding operations. Answering this will require models that not only reproduce representational geometry but also learn it.

1 Neuron and network hyperparameters are provided in the accompanying code repository (URL withheld for anonymous review)

2 While we only include slot-filler vectors constructed with circular convolution here, this factorized property should hold for slot-filler vectors constructed with VTB or tensor products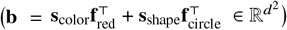.

